# Select Gut Bacteria Promote Mosquito Larval Growth by Contributing Protease Activity and Optimal Ionic Microenvironment for Protein Digestion

**DOI:** 10.1101/2024.02.01.578530

**Authors:** Jhuma Samanta, Sibnarayan Datta

## Abstract

The unique pH profile of mosquito larval midgut including the highly alkaline anterior midgut (AMG) and its significance in digestive physiology has been known for a long time. More recently, metagenomic association studies have shown that diverse assemblage of bacteria (microbiota) present in the gut influence larval nutrition, growth, development, and several other physiological processes. However, the precise functional mechanisms of these host-microbe interactions remain poorly understood. In this research, we studied the functional significance of gut associated bacteria in larval growth utilizing *Aedes albopictus* axenic/ gnotobiotic model. Results of our study demonstrate that, for AMG alkalinization, concurrent presence of Carbonic Anhydrase (CA) activity and ‘*live*’ bacteria is essential and that the later could not be substituted by either cell-free bacterial culture supernatant, or bacterial cell lysate. Further, we found that the gut pH profile and presence of proteolytic activity could be observed in larvae colonized only with select members of the microbiota that are capable of secreting extracellular protease. Collectively, our findings suggest a novel and interesting evolutionary mechanism of host-microbe interaction, in which select members of the gut microbiota promote larval nutrition, growth and development by providing protease and creating ionic microenvironment optimal for its activity.

## INTRODUCTION

Insects are one of the most diverse and adapted life forms on the earth. Certain insects have evolved strategies for maximal exploitation of available resources through switching of ecological niche and feeding habits at various stages of their life cycle (Douglas, 2011; Gullan & Cranston, 2014). The natural ability of insects to feed upon and digest a wide variety of substrates available in different habitats has been linked to the unique physiology and microbiology of its gut; and thus, insect gut has been a subject of intense research for long (Serrato-Salas & Gendrin, 2023; Schmidt & Engel., 2021; Martinson & Strand, 2021; Douglas, 2015). According to feeding habits, gut milieu in herbivorous insects is generally more alkaline as compared to carnivorous insects (Harrison, 2001; Yonge, 1937). Surprisingly, in larvae of certain endopterygote insects belonging to the orders Diptera, Lepidoptera, Coleoptera and Trichoptera, gut pH levels may reach exceptionally high (pH 9.0-12.0), while, extremely low pH have been documented in several others (Gullan & Cranston, 2014; Clark, 1999; Dow, 1984; Berenbaum, 1980; Dadd et al., 1975; Waterhouse, 1949). Like all other organisms, acid-base homeostasis in insects is indispensable for various biochemical and physiological processes including enzyme-substrate reactions, digestion, nutrition, membrane transport of nutrients, vitellogenesis, secretion and functioning of hormone, excretion of nitrogen compounds, pesticide metabolism, pathogen infection etc., thereby disturbance in acid-base homeostasis severely affects survival, growth, development, and vector competence etc. (Harrison, 2001).

Among the other insects, mosquitoes have remained at the centerstage of vector research, owing to their significance in public health. Larval stages have been extensively studied, as a potential target for vector-control strategies, which has led to revelation of several intriguing physiological characteristics of its gut (Dow, 1984; Dadd, 1975; Yonge, 1937). Mosquito larvae belonging to various genera display a wide longitudinal pH gradient profile within a short span of the gut, that too without any morphological barrier (Boudko et al., 2001; Dadd, 1975). Mosquito larval midgut includes an extremely alkaline (pH∼10.0-12.0) anterior midgut (AMG), immediately flanked by neutral/ near-neutral gastric cecum (GC) and posterior midgut (PMG), on either side [**Figure 1**]. The level of alkalinity seen in the AMG of these larvae are rarely met in any other living organism and this gastric arrangement is remarkably contradictory to other metazoans, where optimal digestion initiates at an acidic to highly acidic environment. The characteristic gut pH gradient profile in mosquitoes (also in silkworms) has been shown to provide suitable ionic conditions for sequential functioning of digestive enzymes (with different pH optima), necessary for efficient digestion of complex feed materials (Douglas, 2015; Gullan & Cranston, 2014; Berenbaum, 1980; Dadd, 1975; Horie et al., 1963). Recently, a group of researchers have come up with a novel idea of utilizing this unique pH profile for designing and developing novel mosquito larvicidal molecules (protected triazabutadienes) that are activated (produce protein-reactive aryl diazonium ions) only upon exposure to the high pH environment of the mosquito midgut, thereby reducing the chances of effects on off-target organisms (Guzmán et al., 2023). Development and successful application of such vector control strategies require a detailed understanding of the factors that modulate the unique ionic microenvironment of the larval gut.

**Figure 1.**
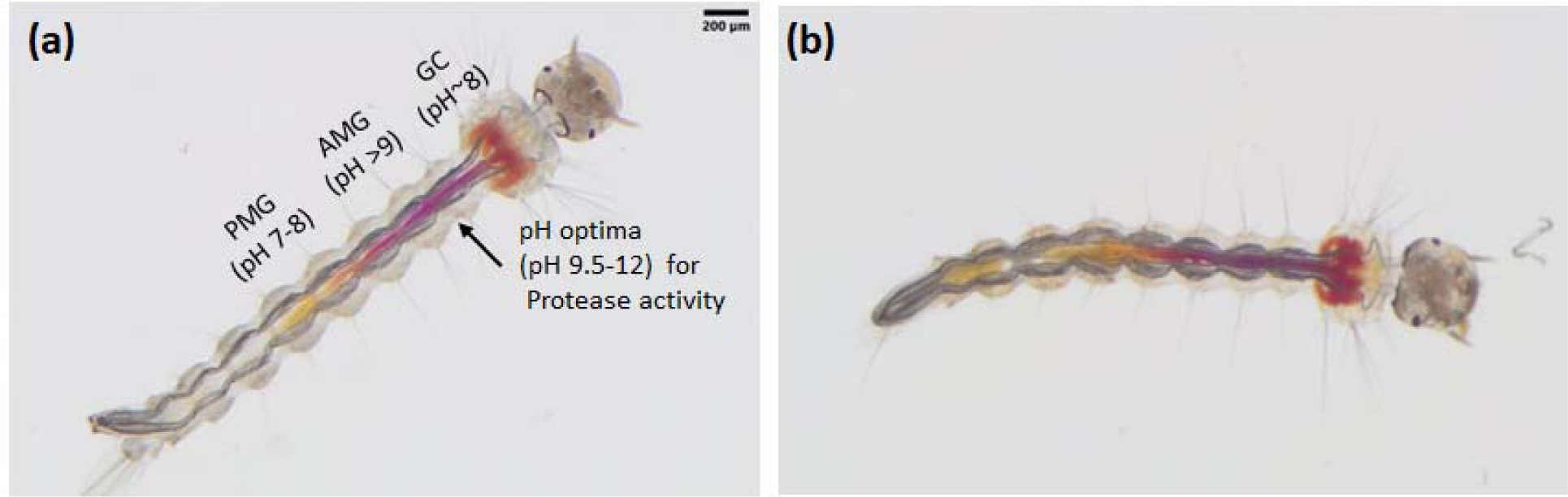
Characteristic gut pH profile as seen in (a) monoculture (*Bacillus cereus*) associated gnotobiotic and (b) conventionally reared *Ae. albopictus* larva, fed with Phenol Red pH indicator dye. Significance of the highly alkaline anterior midgut (AMG) in proteolytic activity is shown following Dadd (1975) in panel (a).

It is also well known for long that mosquito larvae harbor diverse microbiota within their gut, which essentially modulate a wide variety of phenotypes that are associated with the evolutionary success of the host (Schmidt & Engel., 2021; Gao et al., 2019; Strand, 2018; Douglas, 2015; Chao & Wistreich, 1960; Buddington, 1941; Yonge, 1937). Although, an imperative and positive role of gut bacteria in larval survival, growth and development is principally accepted, exclusive requirement of ‘*live*’ bacteria for implementation of such functions has been seriously debated (Schmidt & Engel., 2021; Steven et al., 2021; Coon et al., 2020; Correa et al., 2018; Strand, 2018; Coon et al., 2017; Coon et al., 2014). While a group of researchers claim that ‘*live*’ bacterial signals larval growth and development, others have sternly challenged the proposition (Correa et al., 2018; Valanzia et al., 2018a, b; Coon et al., 2017). However, based upon extensive review of literature, microbiota has been suggested to carry out at least two core functions, nutrition and defense (Serrato-Salas & Gendrin, 2023; Douglas, 2015; Buddington, 1973; Dadd, 1973), both of which are supposed to require ‘*live*’ microbiota.

Nevertheless, comparative studies conducted on axenic and gnotobiotic cultures have demonstrated that mosquito larvae grown under bacteria-free or antibiotic treated conditions along with nutritionally complete diet, may survive for limited periods, but growth and development in such larvae is either stalled at an early stage or is severely delayed; whereas several bacterial species of the gut microbial community or non-community members could individually rescue normal growth and development of axenic larvae, even without additional of nutrient supplements (Wu et al., 2023; Romoli et al., 2023; Martinson & Strand, 2021; Steven et al., 2021; Valzania et al., 2018a, b; Correa et al., 2018; Strand, 2018; Coon et al., 2017; Coon et al., 2014; Chouaia et al., 2012; Buddington, 1941). These observations clearly suggested that larval digestion, growth, and development, are tightly regulated through an evolutionary conserved mechanism, essentially involving gut-bacteria (Schmidt & Engel., 2021; Strand, 2018). Surprisingly however, in the context of exponentially growing volumes of scientific literature on genetic diversity of gut microbiota and its association with diverse phenotypic consequences, information available on mosquito-gut microbiota interaction mechanisms is extremely scanty. Considering the accumulating evidence on the role of larval gut microbiota in nutrition, growth, development, and their carryover effects (in reproduction, fecundity, vector competence etc.) during the subsequent adult stages, elucidation of functional significance of mosquito-gut microbiota interaction in larval physiology and metabolism are essential for controlled manipulation of gut associated microbiota as a potential strategy for vector-control (Harrison et al., 2023; Arellano et al., 2023; Giraud et al., 2022; Gomez et al., 2022; Cansado-Utrilla et al., 2021; Gao et al., 2020; Huang et al., 2020; Scolari et al., 2019).

Incidentally, while studying the effects of bacteriophage mediated manipulation of gut bacteria on gut physiology in *Aedes albopictus* larvae (colonized with *Bacillus cereus*), we observed that the characteristic pH profile in larval gut is eliminated after treatment with highly lytic bacteriophages against *B. cereus*, sharply contrasting the intact pH profile in mock treated (no bacteriophage) larvae [**Figure 2**]. Repeated and reproducible observation of this phenomenon led us to hypothesize a direct role of gut bacteria in larval gut physiology. Subsequently, we tested this hypothesis using *Ae. albopictus* axenic/ gnotobiotic experimental culture model, which resulted in revelation of an interesting mechanism of host-microbe association in the larval gut.

**Figure 2.**
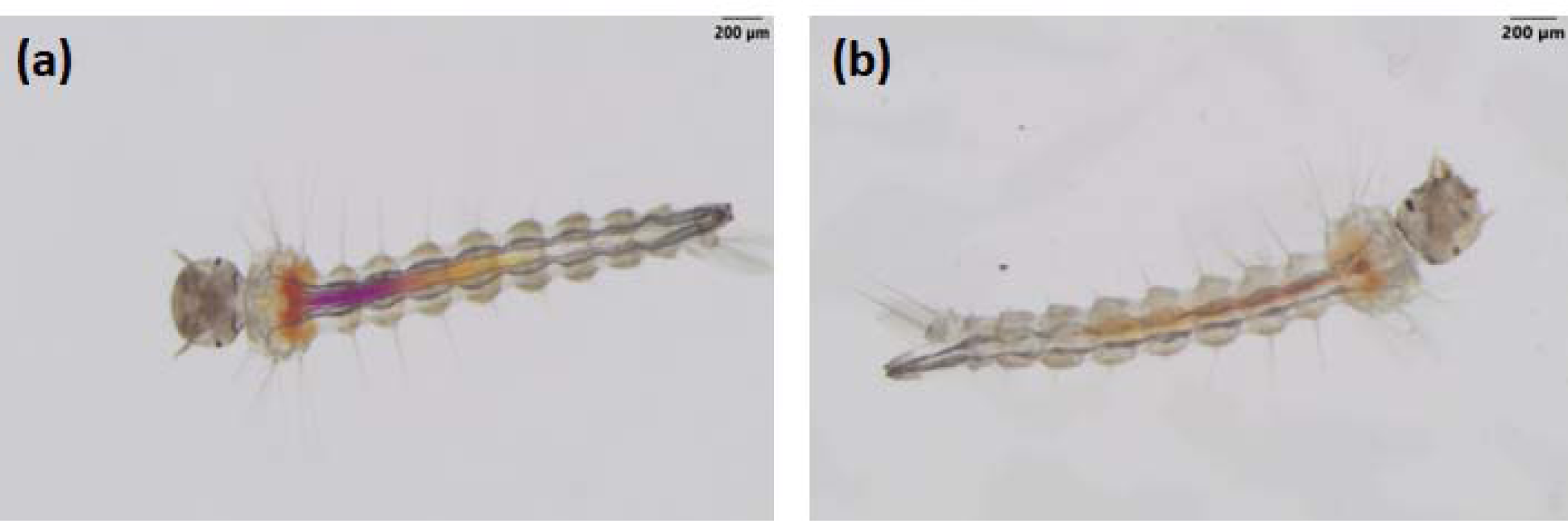
Gut pH profiles in (a) mock treated (no bacteriophage) mono-associated gnotobiotic larvae; (b) bacteriophage treated gnotobiotic larvae (*Bacillus cereus*).

Results of experiments conducted in the present research work revealed that select gut bacteria are directly associated with establishment and maintenance of the gut pH profile along with secretion of protease, essential for digestion and nutrition, thereby promoting larval growth, and development. Through this communication, we report our findings, and discuss their significance under the light of available literature.

## MATERIALS & METHODS

### Generation of axenic, gnotobiotic larvae and gut pH visualization

Axenic larvae were generated by surface sterilizing *Ae. albopictus* eggs, following an earlier protocol (Romoli & Gendrin, 2020) with slight modifications. The sterility of axenic larvae was verified by culturing specimen larval homogenates on LA plates for 48 hrs. A subset of axenic larvae was then used to generate monoculture gnotobiotic larvae by colonization with a strain of *Bacillus cereus* (GenBank Accession No. OR513030), which we previously isolated from laboratory reared *Ae. albopictus* larvae. For initial experiments, this bacterial species was selected as it has been reported to colonize the larval gut of different mosquito species stably and harmlessly for longer periods without interfering with their growth and development (Luxananil et al., 2001). Selective colonization of bacteria in the gut was carried out by adding fresh bacterial culture (@ 10^6^ CFU mL^-1^) to the axenic larval culture plate-wells. After three days post colonization, larvae were collected, washed thoroughly but gently with sterile water, and transferred to fresh plate-wells holding sterile water. Stable bacterial colonization was confirmed by culturing homogenates from thoroughly washed specimen gnotobiotic larval on LA plates for 48 hrs, followed by morphological examination of the colonies.

All larval culture experiments were carried out at 27±2°C with 80±10% RH. All microbiological manipulations were performed inside class II (A2) biological safety cabinet, and standard microbiological laboratory procedures and precautions were followed strictly.

Larval gut pH profile was visualized using 0.1 % w/v phenol red (PR) pH indicator dye fed through addition in the holding media, as described previously (Overend et al., 2016; Boudko et al., 2001; Dadd, 1975), with minor modifications. Larvae were allowed to ingest the dye for at least 15 mins before visualization.

### Study of gut pH profile in experimentally treated larvae

Gut pH profiles in axenic, gnotobiotic (colonized by *B. cereus*), antibiotic (Gentamycin @ 20 µg mL^-1^) treated gnotobiotic, and Acetazolamide (@ 100 µM) treated gnotobiotic larvae were observed and documented under a stereomicroscope (SZX16, Olympus, Tokyo, Japan). Gut pH profiles were visualized, following pH indicator dye method described above. Larvae from *Ae. albopictus* culture, conventionally reared in open tray under standard laboratory conditions were included as natural controls.

To further verify if component(s) from bacterial cultures (cell free supernatant/lysate) could substitute the requirement of ‘*live*’ bacteria for establishment of gut pH profile, we studied the gut pH profiles in axenic larvae, axenic larvae reared with bacterial cell free culture supernatant (*B. cereus*), axenic larvae reared with bacterial cell lysate (axenic larvae reared with *B. cereus*) and gnotobiotic larvae (axenic larvae reared with live *B. cereus*). Larval length was recorded 4 days post treatment to compare variation in growth rates of larvae from different experimental groups.

### Gut pH profile in gnotobiotic larvae colonized with different bacteria

To examine if any of the members of the gut-microbiota could establish pH gradient profile or only select members have the capability, we studied the gut pH profile in various monoculture gnotobiotic larvae, individually colonized with *B. cereus* (GenBank Accession No. OR513030)*, Aeromonas hydrophila* (OR513036)*, Microbacterium paraoxydans* (OR513035)*, Leucobacter coleopterorum* (OR513031)*, Lysinibacillus fusiformis* (OR513032), all of which were previously isolated from laboratory reared *Ae. albopictus* larvae. Axenic larvae served as control. Larval length was recorded 4 days post colonization to compare the pH profile and variation in growth rates of larvae from different experimental groups.

### Proteolytic activity assay

The presence of proteolytic activity in different monoculture gnotobiotic larvae was qualitatively evaluated by incubating larval homogenates on standard casein agar plates. In brief, equal numbers of larvae (50) from different experimental groups, as mentioned in the preceding section were collected, rinsed gently, homogenized, and clarified by centrifugation at 8,000 *g* for 10 mins at 4°C. The supernatant was then collected and used for casein and starch hydrolysis assays. A volume of 100 µL of each clarified lysate were then added separately to designated casein agar plate (supplemented with Gentamycin @ 20 µg mL^-1^) wells and incubated overnight at 25 °C. Subsequently, zones of hydrolyses were visualized after treating the plates with trichloroacetate and photographed. Area under zones of hydrolysis were calculated with the help of an open-source computer program, ImageJ (Schneider et al., 2012).

To verify the protease secretion capability of the gut associated bacteria included in this study, fresh cultures of each bacterium were streaked on casein agar plates, incubated and zone of hydrolysis visualized as mentioned above.

## RESULTS

### Presence of live bacteria is required for establishment of distinctive pH profile in larval gut

Our results show that, in sharp contrast to the distinctive gut pH profile seen in conventionally reared and gnotobiotic larvae, no such profile was observed in axenic, antibiotic treated, or acetazolamide treated larvae [**Figure 3**]. While antibiotic treated larvae presented the bactericidal effects of gentamicin on gut colonized bacteria, acetazolamide treatment presented the effects of carbonic anhydrase (CA) inhibition in presence of gut bacteria, suggesting concurrent requirement of gut associated bacteria and CA activity for midgut alkalinization.

**Figure 3.**
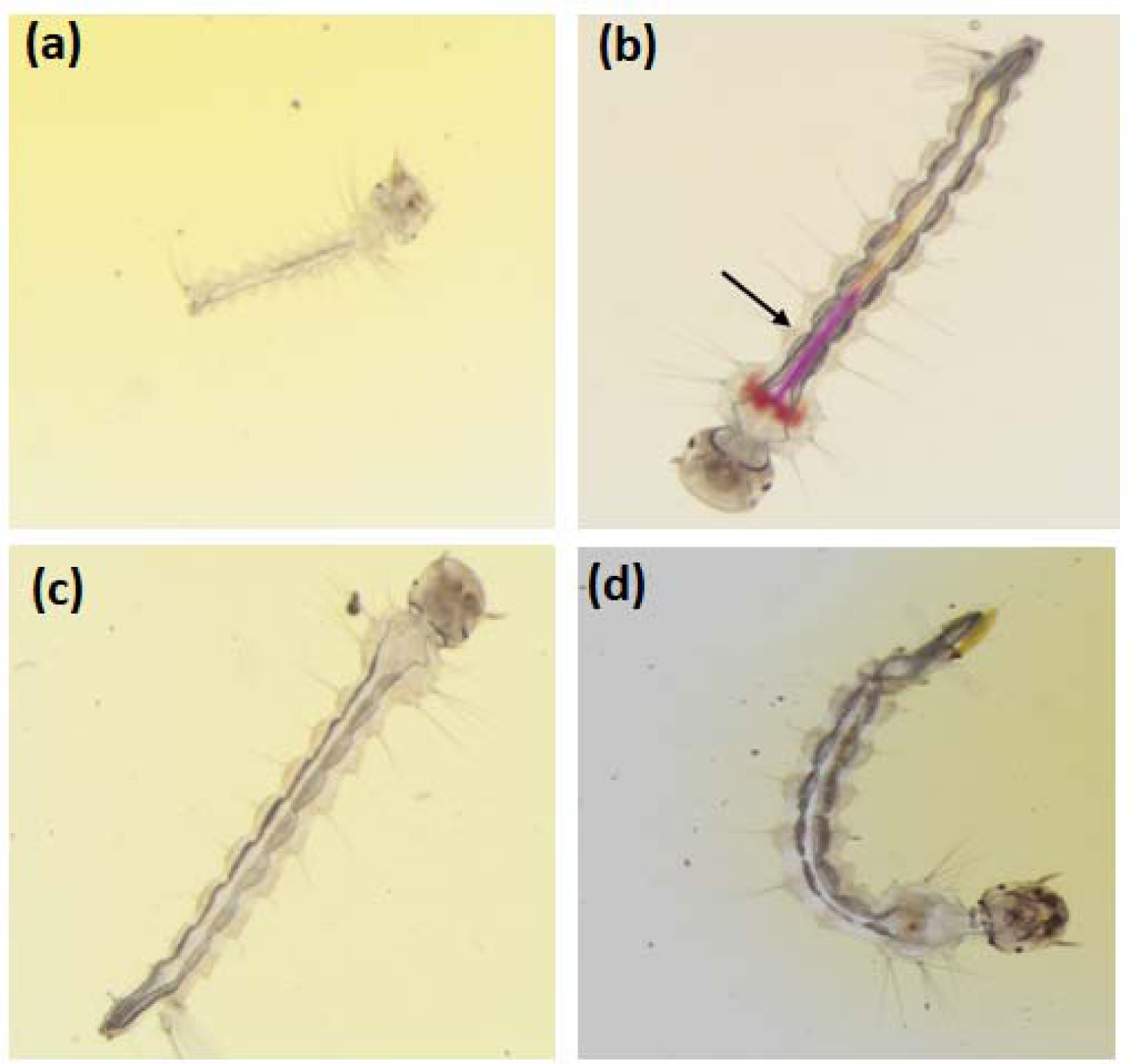
Gut pH profiles in (a) axenic; (b) gnotobiotic; (c) acetazolamide treated gnotobiotic and (d) antibiotic (Gentamycin) treated gnotobiotic larvae (*Bacillus cereus*).

### Continual presence of live bacteria is required for maintenance of distinctive pH profile in larval gut

When gnotobiotic larvae displaying a well-established gut pH profile were subjected to antibiotic treatment, the distinctive pH profile was found to be completely abolished. This observation suggested that establishment and maintenance of gut pH profile by gut bacteria is not a typical ‘*hit and run*’ phenomenon, rather it requires continual presence of bacteria [**Figure 4**].

**Figure 4.**
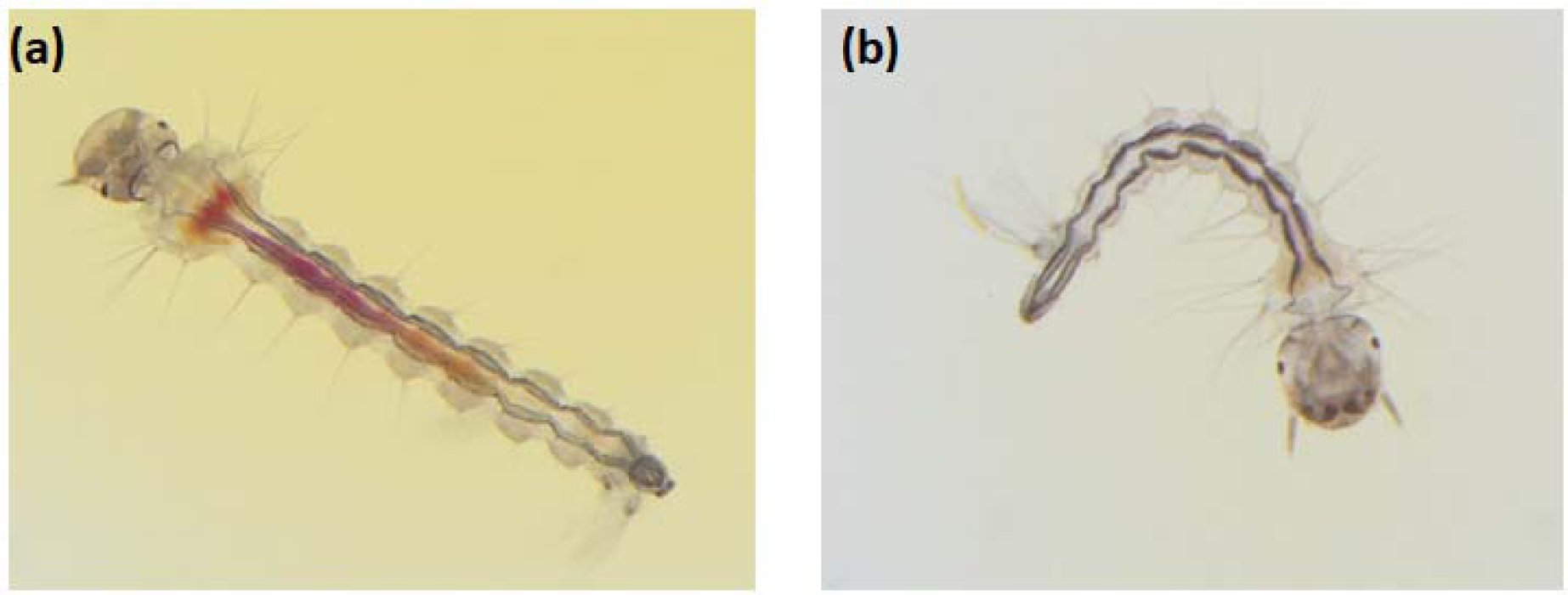
Gut pH profiles in (a) gnotobiotic larvae (before antibiotic treatment); (b) gnotobiotic larvae 24 hrs after treatment with antibiotic (Gentamycin).

### Only live bacteria can establish and maintain larval gut pH and is associated with larval growth

Our results with larvae grown with live bacteria, cell-free bacterial culture supernatant and bacterial cell lysates demonstrate that the distinctive pH profile is established only in larvae reared with live bacteria, but neither in larvae reared with bacterial culture supernatant nor with bacterial cell lysate [**Figure 5 a-d**]. Further, growth of larvae reared with live bacteria having distinctive pH profile was significantly higher (P<0.001) than the growth of larvae reared under axenic condition or with bacterial cell lysate or bacterial culture supernatant [**Figur**e 5 e].

**Figure 5.**
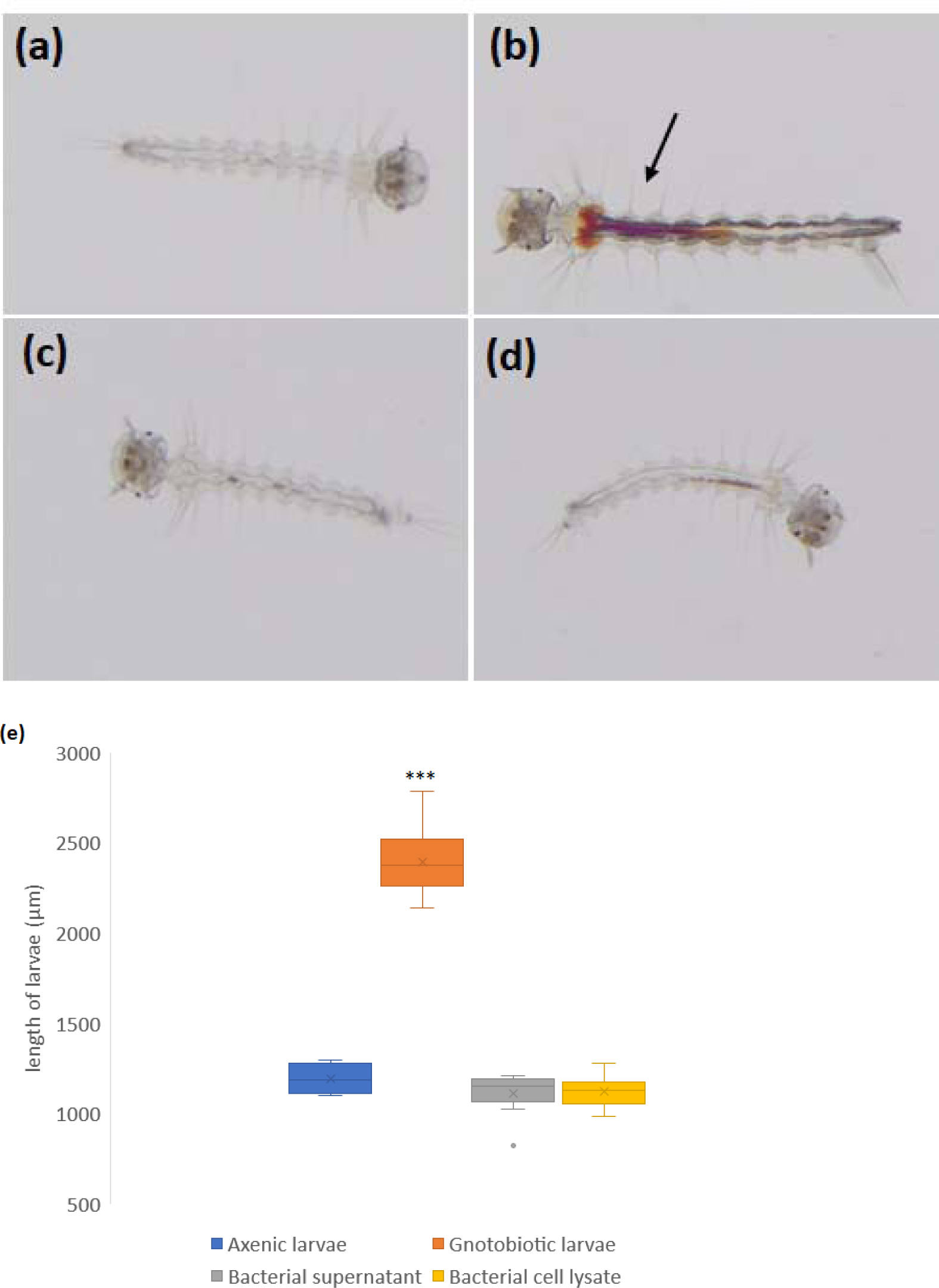
Gut pH profiles in (a) axenic larvae; (b) axenic larvae grown in presence of live bacteria (gnotobiotic, *Bacillus cereus*); (c) axenic larvae grown in presence of bacterial cell-free culture supernatant (*Bacillus cereus*); (d) axenic larvae grown in presence of cultured bacterial cell lysate (*Bacillus cereus*); (e) graph showing comparison of larval length from each experimental group (n≥10 in each group).

### Only select members of the gut microbiota can establish distinctive gut pH profile, modulate proteolytic activity and promote larval growth

We observed that among the members of the gut microbiota included in the present study, bacteria from 2 different families-*Bacillaceae (B. cereus* and *L. fusiformis)*, *Aeromonadaceae (A. hydrophila*) could establish the distinctive gut pH profile and promote larval growth, while bacteria from the family *Microbacteriaceae* (*M. paraoxydans, L. coleopterorum*) could not [**Figu**re 6, a-g]. Lengths of gnotobiotic larvae reared with *B. cereus, L. fusiformis* or *A. hydrophila* were significantly higher than the axenic larvae or gnotobiotic larvae reared with *M. paraoxydans, L. coleopterorum*.

**Figure 6.**
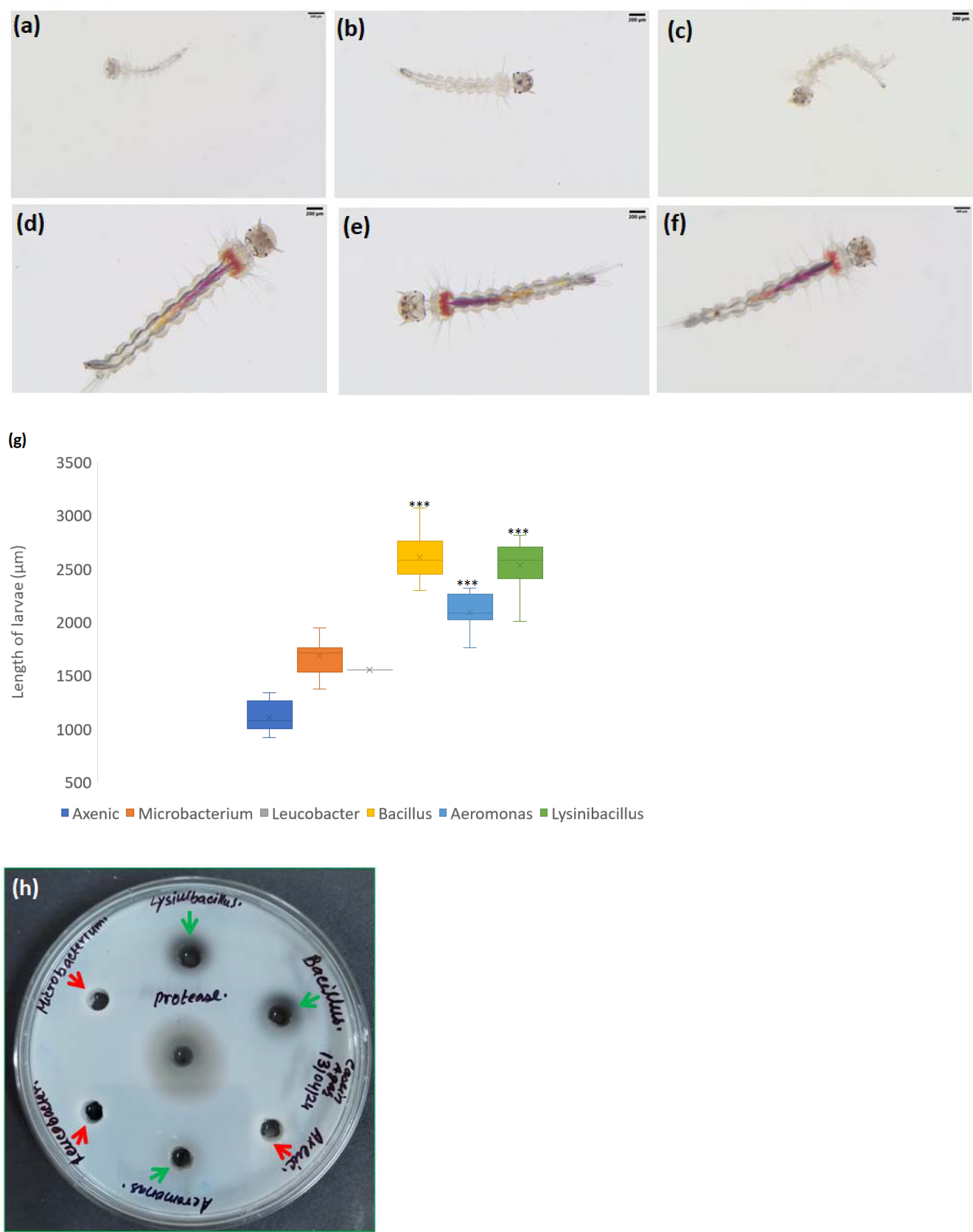
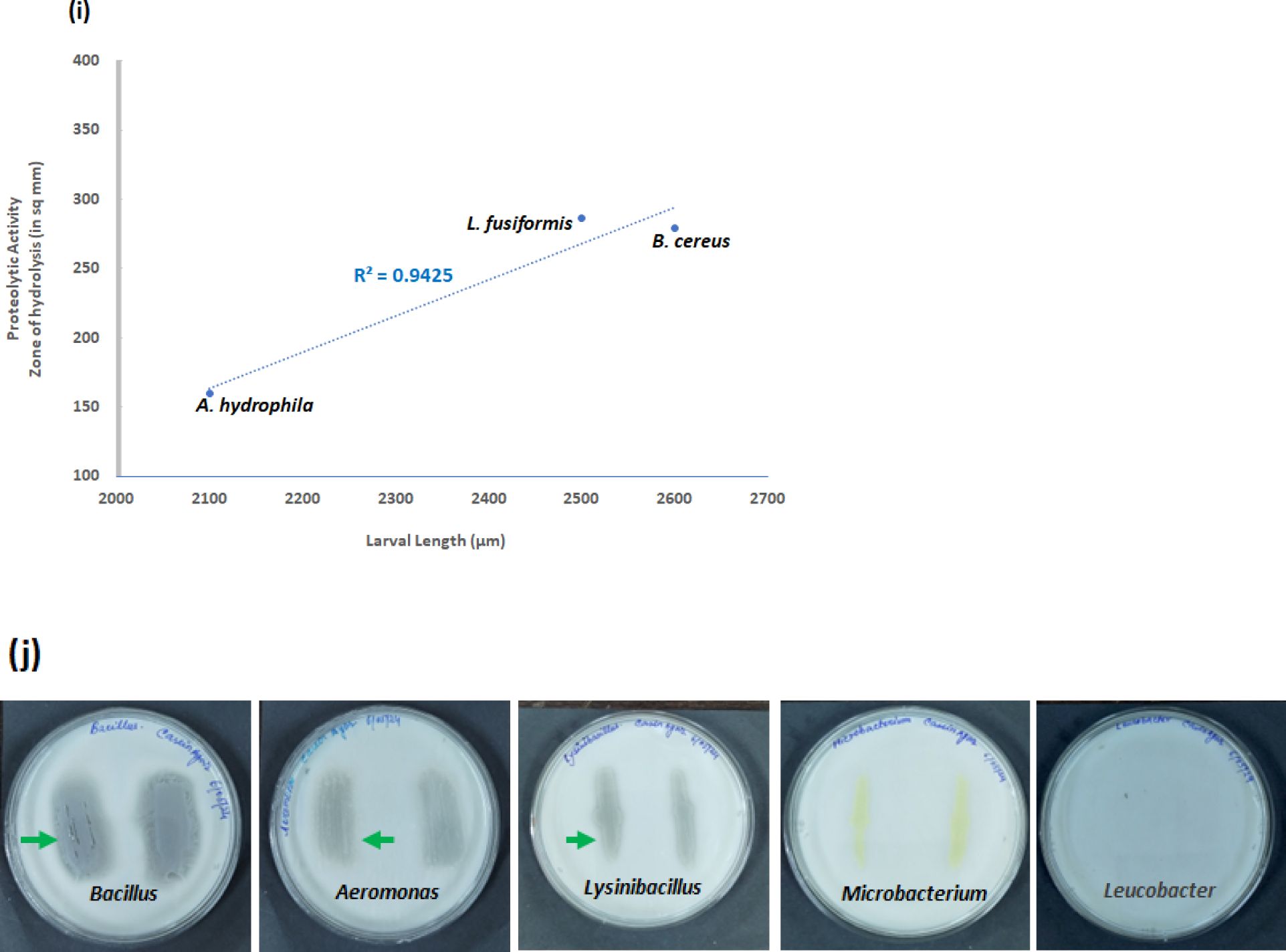
Gut pH profiles in (a) axenic larvae and monoculture gnotobiotic larvae, individually colonized with different bacteria-(b) *M. paraoxydans;* (c) *L. coleopterorum*; (d) *B. cereus*; (e) *A. hydrophila*; (f) *L. fusiformis*; (g) graph showing comparison of larval length as an indicator of larval growth from each experimental group; (h) Casein agar plate showing proteolytic activity in gnotobiotic larvae reared with different bacteria. Lysates showing proteolytic activity are indicated by green arrow heads, while lysates showing no proteolytic activity are shown by red arrow heads; (i) Scatterplot showing correlation between proteolytic activity and larval length; (j) Casein agar plates showing secretory protease activity (indicated by green arrow heads) associated with different gut bacterial culture studied in this research.

Detection of proteolytic activities in lysates from different groups of gnotobiotic larvae corresponded with the presence of distinctive gut pH profile and larval growth [**Figure 6 h**]. Markedly, gnotobiotic larvae reared with *B. cereus* and *L. fusiformis* showed higher levels of proteolytic activity and growth, as compared to larvae reared with *A. hydrophila* [**Figure 6, g, h**]. When larval growth data and proteolytic activity from these 3 groups (*B. cereus, L. fusiformis* and *A. hydrophila*) of gnotobiotic larvae were analyzed, a highly positive correlation between proteolytic activity and larval length was recorded (R^2^=0.9425) [**Figure 6, i**]. No proteolytic activity was observed in lysates from axenic larvae or gnotobiotic larvae reared with *M. paraoxydans, L. coleopterorum* [**Figure 6, h**].

Furthermore, when each of these 5 bacteria were grown individually on casein agar plates, *B. cereus, L. fusiformis* and *A. hydrophila* showed extracellular protease activity, while no proteolytic activity was observed on casein plates inoculated with *M. paraoxydans*. On the other hand, *L. coleopterorum* did not even grow on the casein plates [**Figure 6, j**].

## DISCUSSION

Results of our experiments showing comparable gut pH profiles in conventionally grown larvae (harboring a complex microbial community) and in select monoculture gnotobiotic larvae (*Bacillus, Aeromonas, Lysinibacillus*) suggests a broadly taxon-specific host-microbe collaboration in the present scenario. Our finding of a functional pH gradient in monoculture gnotobiotic larvae colonized with *Aeromonas*, but not in larvae colonized with *Microbacterium* or *Leucobacter*, strikingly agrees with previous rescue, growth and development studies showing that larvae reared with *Aeromonas* survive, grow, and develop normally, while those with *Microbacterium* do not (Coon et al., 2014). Our results also provide an explanation to this gut bacteria genera/species specific capability of rescuing axenic larvae and supporting their growth and development.

In this study, non-establishment of pH profile in acetazolamide treated gnotobiotic larvae also reconciles with earlier electrophysiological and biochemical studies, reiterating an intrinsic role of CA mediated ionic homoeostatic mechanisms in AMG alkalinization and in survival and growth of larvae (Francis et al., 2016; Linser et al., 2009; Corena et al., 2004). Apparently, the conventional notion that colonization or decolonization of microbiota takes place as a function of gut pH (Schmidt & Engel., 2021; Coon et al., 2017; Dillion & Dillion, 2004) indicates a role of larval CA activity, but we could not find any compelling evidence for a dominant role of larval CA in midgut alkalinization in the relevant literature. Notably, in earlier localization studies, patterns of CA expression in host tissues did not correlate with the gut alkalinization profile (very high CA expression in near-neutral GC and PMG region, while no or very low expression in highly alkaline AMG), based on which, involvement of ‘soluble extracellular CAs’ in AMG alkalinization was hypothesized (Linser et al., 2009; Onken et al., 2008; Corena et al., 2005; Seron et al., 2004; Corena et al., 2002). Likewise, previous studies showing the capability of live intact larvae in generating high luminal pH, while dramatic loss of this capability in dissected out, *in vitro* bathed, perfused gut preparations seem to support an obvious role of the hypothesized ‘soluble extracellular CA’ as the principal factors involved in gut alkalinization (Onken et al., 2008; Onken et al., 2004). Even though the exact microbial status of the larvae in aforesaid studies were not specifically stated in the literature, considering their reported conventional rearing conditions, we assume that all these larvae carried conventional microbiota. Indeed, when we reviewed these observations from the perspective of our present findings, there appears an intricate association between the suggested ‘soluble extracellular CA’ and the gut microbiota.

To the best of our knowledge, bacteria specific CA activity has not been formally characterized in the mosquito larval gut, till date. However, several studies describing secretory CA activity in the gut lumen, presence of extracellular CA activity especially in the ectoperitrophic fluid and luminal detection of CA belonging to certain classes (e.g., α & β classes of CA that are known to be encoded by various prokaryotes too) do appear to imply a prominent role of gut bacteria secreted CA in luminal alkalinization (Vullo et al., 2018; Syrjanen et al., 2010; Smith et al., 2007; Fisher et al., 2006; Corena et al., 2005; Seron et al., 2004; Corena et al., 2002). Such a possibility is also firmly supported by strong expression of immunologically CA-like gene products by certain bacterial genera (e.g. *Pseudomonas, Serratia and Proteus*), frequently associated with mosquito microbiota (Scolari et a., 2019; Nafi et al., 1990). Notably, a recent study on another Dipteran larvae (*Drosophila*) has revealed an interesting mechanism of modulation of gastrointestinal contractions mediated through acetylcholine secreted by a gut associated bacterium, *Lactiplantibacillus plantarum* in a species-specific manner (Fujita et al., 2024), which reiterates a direct and species-specific role of gut associated bacteria in various aspects of larval physiology.

On the other hand, the fact that certain Dipteran and Lepidopteran insects host a remarkably alkaline midgut environment, which provides functionally optimal ionic conditions for various digestive enzymic activities related to nutrition, growth and development, especially the highly alkaline protease activity, has been revealed in early studied (Eguchi & Iwamoto, 1976; Dadd, 1975; Yang & Davis, 1972; Gooding, 1966; Horie et al., 1963). Through exhaustive physiological and biochemical studies on larvae belonging to distinct mosquito genera (*Aedes, Culex, Anopheles*), Dadd (1975) demonstrated that the characteristic gut pH profile remains intact throughout different instars, irrespective of presence of food in gut and that pH optima of important digestive enzyme complexes are remarkably similar across diverse genera, suggesting gut pH profile as a distinctive evolutionarily conserved taxonomical feature (Dadd, 1975). Further, based on the observed range of pH optima of amylase (7.0-9.0) and protease (9.5-12.0), he concluded that protein digestion is strictly restricted to the highly alkaline AMG milieu, while starch digestion is limited to the neutral to moderately alkaline PMG (Dadd, 1975). These observations clearly underline the crucial role of the distinct larval gut pH profile and an alkaline protease in dissociation and coagulation of ingested protein complexes, regulation of digestion reactions, neutralizing toxic materials, all of which contribute towards survival, growth, and development of the larvae (Gullan & Cranston, 2014; Berenbaum, 1980; Dadd, 1975; House, 1974). In this background, our present findings add another level of complexity by proposing a novel facet of host-microbe interaction in larval digestive physiology, where members of the gut associated microbiota belonging to certain genera/ species mediate larval growth and development through establishment of gut ionic microenvironment optimal for important enzymic activities.

In the context of highly alkaline AMG, several studies have documented the dominance of diverse serine type proteases in mosquito larval gut (Saboia-Vahia et al., 2013; Kunz, 1978; Dadd, 1975; Yang & Davis, 1971). Even though the possibility that some of these proteases may derive from the gut microbiota was suggested long back (Kunz, 1978), the connection between gut microbiota and proteases in mosquito larvae has remained understudied, as compared to other Lepidopteran larvae. Nevertheless, we could find ample suggestions of significant role of gut bacteria produced proteolytic activity in mosquito larvae from previous studies that show strong interference in mosquito larval protein degradation capabilities, survival, growth and development following treatment with protease inhibitors, bactericidal agents, or antibiotics (Sajna et al., 2019; Pontual et al., 2014; Patil et al., 2013). Analogous inhibition of hemolytic/ proteolytic and other enzymic activities after treatment with antibiotic/ antiseptic has also been shown in several other insects, including bollworm, silkworm, velvetbean caterpillar, adult mosquitoes, and arachnids (O Gaio et al., 2011; Regode et al., 2016; Visotto et al., 2009; Yang & Davis, 1972; Eguchi & Iwamoto, 1976; Gooding, 1966; Horie et al., 1963; Terzian, 1958). Interestingly, in early studies on mosquito larvae and silkworm, drastic decline in gross proteolytic activity has been documented during or post larval-pupal ecdysis phase, which strikingly coincide with radical reorganization of gut structures causing major turnover or complete removal of the gut microbiota during metamorphosis in these insects (Manthey et al., 2022; Eguchi & Iwamoto, 1976; Yang & Davis, 1971; Horie et al., 1963). Apart from proteolytic activity, gut microbes are also known to provide various other nutrients and enzymes that help the insect hosts in digesting complex substrates (Douglas, 2009).

Agreeing and furthering the previous studies, our present results demonstrate that select members (*Bacillus, Lysinibacillus* and *Aeromonas* in this study) of the gut microbiota are directly connected to the presence of protease activities in the larval gut. These findings agree with a previous study on Lepidopteran larvae, showing that among the diversity of bacteria present in the gut, not all, but a select extracellular protease producing gut bacteria contribute to the proteolytic activity in the larval gut (Regode et al., 2016). Additionally, our data also suggests that the level of larval protease activity differs between genera of colonized bacteria that in turn correlates well with larval growth. Similar gut protease activity dependent variation in growth and development has recently been described in diamondback moth larvae (Zhao et al., 2019), which also attests to the evolutionary significance of this host-microbe interaction mechanism.

## CONCLUSIONS

In conclusion, results of our present study challenge the conventional notion that colonization by microbiota is a function of gut pH and show that this host-microbe association works the other way, where certain members of the gut microbiota establish and sustain the larval gut pH. Further, based upon our findings, we propose a novel and interesting evolutionary mechanism of host-microbe interaction, wherein certain members of the gut microbiota promote larval nutrition, growth and development by providing essential digestive enzyme (protease) and creating ionic microenvironment suitable for optimal activity of the enzyme. We firmly believe that our findings will pave new avenues for future research towards better understanding of the role of gut microbiota in vector biology and in developing efficient, eco-friendly vector-control strategies. Additionally, it would be interesting to examine if similar mechanisms also work for other animals including humans.

## ACKNOWLEDGEMENTS

This work was funded by intramural grants and research fellowship (JS) from Defence Research & Development Organization (DRDO). Authors thank Dr. Bipul Rabha, Dr. Diganta Goswami and Dr. Rajan Pilakandy for technical help. Authors acknowledge the support and encouragement of Dr. Vanlalhmuaka and Dr. Dev Vrat Kamboj. Authors are thankful to Director, Defence Research Laboratory, Tezpur for granting permission to publish the study data (manuscript reference number DRL/EBM/01/2024).

## AUTHOR CONTRIBUTIONS

JS and SD conceptualized the study. JS performed all the experiments. JS and SD analyzed data, wrote, reviewed, and edited the manuscript.

